# Development and evaluation of a reverse transcription loop-mediated isothermal amplification assay for rapid typing of serotype O foot-and-mouth disease virus in endemic regions of Tanzania

**DOI:** 10.1101/2023.04.07.536083

**Authors:** Sarah N. Mueni, Sengiyumva E. Kandusi, Emma P. Njau, Christopher J. Kasanga

## Abstract

Foot-and-mouth disease (FMD) caused by FMD virus, is a highly infectious viral disease affecting livestock. Diagnosis plays a vital role in disease control and management during disease outbreaks. Available serotyping approaches are costly, time-consuming, requires complex equipment and trained personnel, warranting development of a diagnostic method that addresses this gap. In this retrospective study, a reverse transcription loop-mediated isothermal amplification (RT-LAMP) assay for the detection of serotype O FMDV was developed and evaluated using forty-three FMDV isolates from cattle. The specificity of the assay was tested against other FMDV isolates (serotypes A, SAT 1 and SAT 3). Only FMDV serotype O samples could be amplified within 26 min with an anneal derivative temperature of 87.02 degree celsius. Additionally, the assay amplified the viral protein 1 (*VP1*) gene with a detection limit of 0.0378 ng/μl. This molecular diagnostic approach has potential future application in improving FMDV surveillance as it provides baseline information for controlling FMD outbreaks in Tanzania and other Eastern and Southern African countries.

## 1. Introduction

Foot-and-mouth disease (FMD) is a transboundary animal disease (TAD), affecting all cloven-hoofed animals and causes economic strain on livestock production due to trade embargo on livestock and their products (OIE and FAO, 2018).The causative agent is FMD virus (FMDV; genus *Aphthovirus*; family *Picornaviridae*), a non-enveloped, single-stranded, positive-sense RNA virus with 8.5 kb genome with seven immunologically distinct serotypes (O, A, C, Asia 1, [SAT] types 1–3) (Grubman & Baxt, 2004; Jamal & Belsham, 2013). Within each serotype, genotypes and/or topotypes with significant antigenic variability exist and protection from one serotype does not confer immunity against other circulating serotypes (Knowles & Samuel, 2003). The global distribution of FMDV serotypes is uneven, with serotype O and A virus being dominant in sub-Saharan countries (Jamal & Belsham, 2013). serotype O FMDV is further categorized into 11 topotypes: East Africa 1 to 4 (EA 1-4), West Africa (WA), Southeast Asia (SEA), Europe-South America (EURO-SA), Indonesia-1 and −2 (ISA 1-2), Cathay and Middle East-South Asia (ME-SA) (Samuel & Knowles, 2001;Knowles & Samuel, 2003). The high antigenic diversity of FMDV variants makes the disease difficult to control.

Foot-and-mouth disease is endemic in Tanzania, with outbreaks occurring throughout the year in different regions of the country. Serotype O viruses have the widest distribution (Kasanga et al., 2015). Lack of timely diagnosis has been implicated as one of the main hindrances to the control and elimination of FMD. Due to its infectious nature and economic implications, rapid and sensitive diagnostic tests for identifying FMDV serotypes causing the disease outbreak(s) is imperative (Jamal & Belsham, 2013; Kasanga et al., 2014). Molecular assays have been widely employed for the detection of FMDV serotypes, especially real-time reverse transcription PCR (rRT-PCR), which is also recommended by the World Organisation for Animal Health (OIE) (Callahan et al., 2002; OIE, 2018). However, the serotyping is costly, time-consuming, requires complex equipment and trained personnel, hence unsuitable in resource-limited settings of low- and middle-income countries. Hence, alternative molecular technique for detecting FMDV serotypes is warranted to address this gap.

Developed by (Notomi et al., 2000), loop-mediated isothermal amplification (LAMP), is a simple, rapid, sensitive and specific molecular diagnostic tool for detecting viruses of both human and animal origin (Dhama et al 2014; Mori & Notomi, 2009). Reverse transcription loop-mediated isothermal amplification (RT-LAMP) assay for detecting FMDV was first developed by Dukes et al., (2006) followed by several assay development and improvement for detecting FMDV serotypes A, C and O (Chen et al., 2011; Ding et al., 2014; Farooq et al., 2015; Lim et al., 2018; Madhanmohan et al., 2013; Maryam et al., 2017). These assays have been developed specifically for isolates prevalent in China, Pakistan, South Korea, and India respectively. However, the suitability of these assays for the diagnosis of FMDV serotype O circulating in other countries, particularly sub-Saharan Africa has not yet been verified.

The current study aimed to develop a RT-LAMP assay using newly designed primers for the detection of serotype O FMDVs circulating in Tanzania and to verify the suitability of RT-LAMP assay from Pakistan (Maryam et al., 2017) on Tanzania serotype O viruses. This is the first serotype-specific RT-LAMP assay that has been developed in sub-Saharan Africa for diagnosis of FMD. The information gathered from this study is a valuable resource for adoption of appropriate control measures and surveillance of FMD in Tanzania.

## 2. Material and Methods

### 2.1. Study Area

This study was conducted using samples that were previously collected from cattle that showed clinical signs consistent with FMD, from 10 regions of Tanzania (Figure 1 and table 1) and the samples archived in laboratory repository at −80°C.

**Figure 1:**
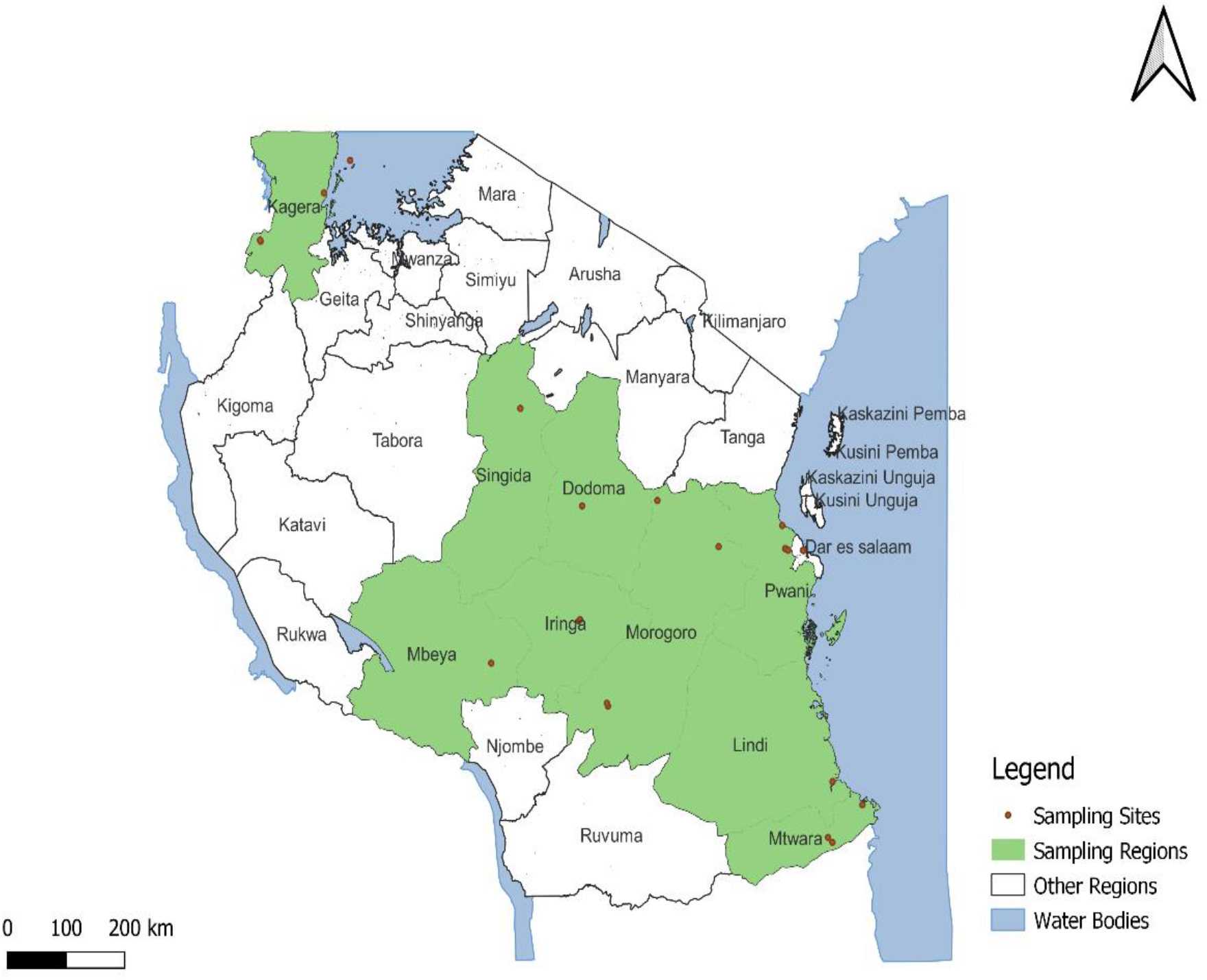
Sampling sites in 10 regions of Tanzania where the FMDV-positive samples originated. Red points denote the location of the sampling sites. The FMDV serotype O-positive samples originated from the regions of Kagera, Morogoro, Pwani, Dar es salaam, Lindi and Mtwara.

**Table 1.**
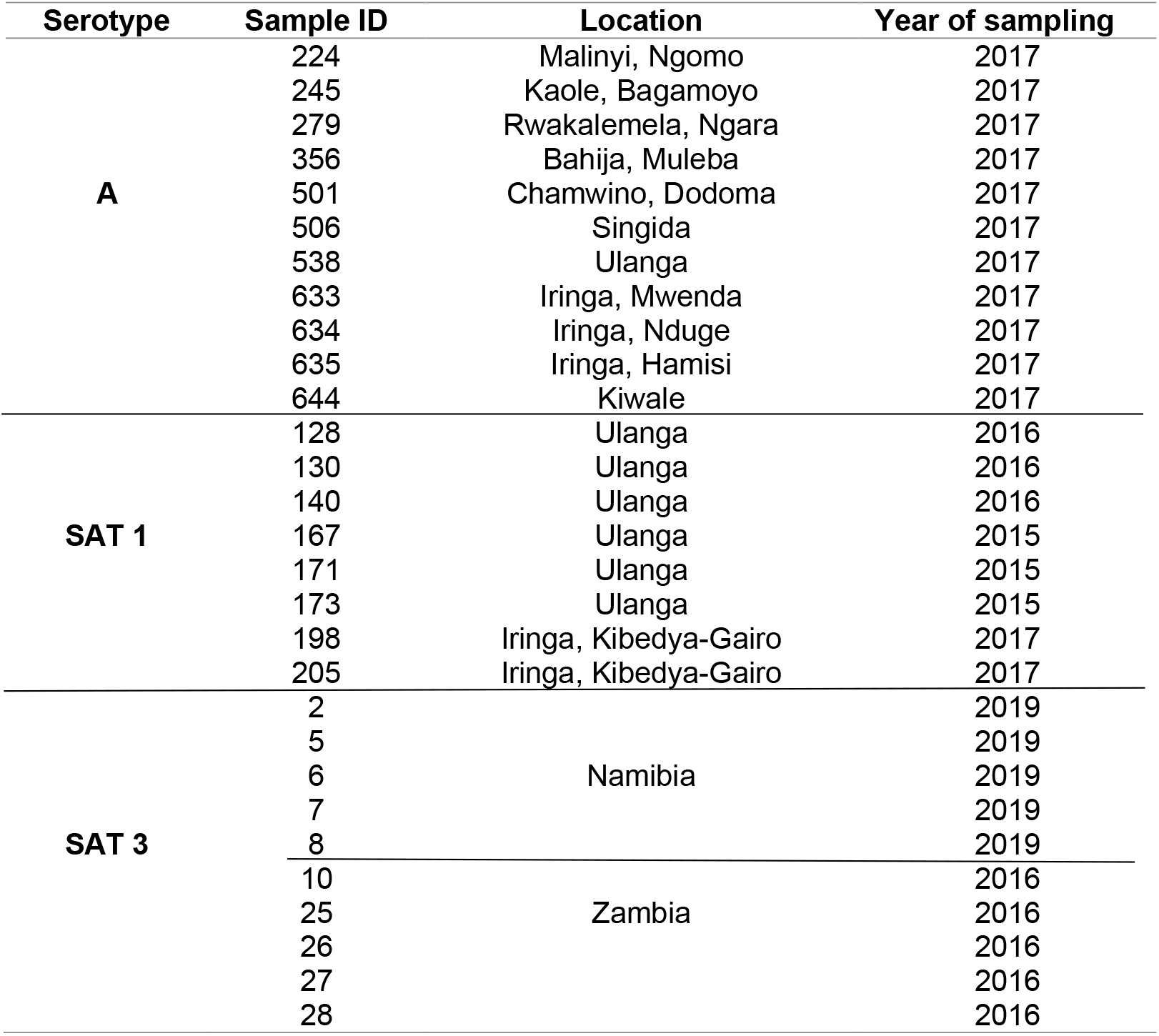
FMDV samples used for evaluation of RT-LAMP assay

### 2.2. FMDV Isolates

Forty-three characterized samples (i.e., previously tested and confirmed to be FMDV-positive using RT-PCR and Sanger sequencing (unpublished data) were used for evaluation of RT-LAMP (n=14 samples; Table 4) and assessing the sensitivity and specificity of the test (n=29 samples; Table 1). Among the 43 samples; 14 serotype O samples were randomly selected for serotyping, and 20 samples for specificity studies (11 serotype A and 9 SAT 1 sample selected from different regions in Tanzania). In addition to the Tanzania samples, 10 SAT 3 samples from Zambia and Namibia were used in specificity study. At the time of study there we no confirmed cases of SAT 2 and SAT 3 in Tanzania (suspected SAT 2 and SAT 3 samples tested negative for viral RNA using RT-PCR).

**Table 2.**
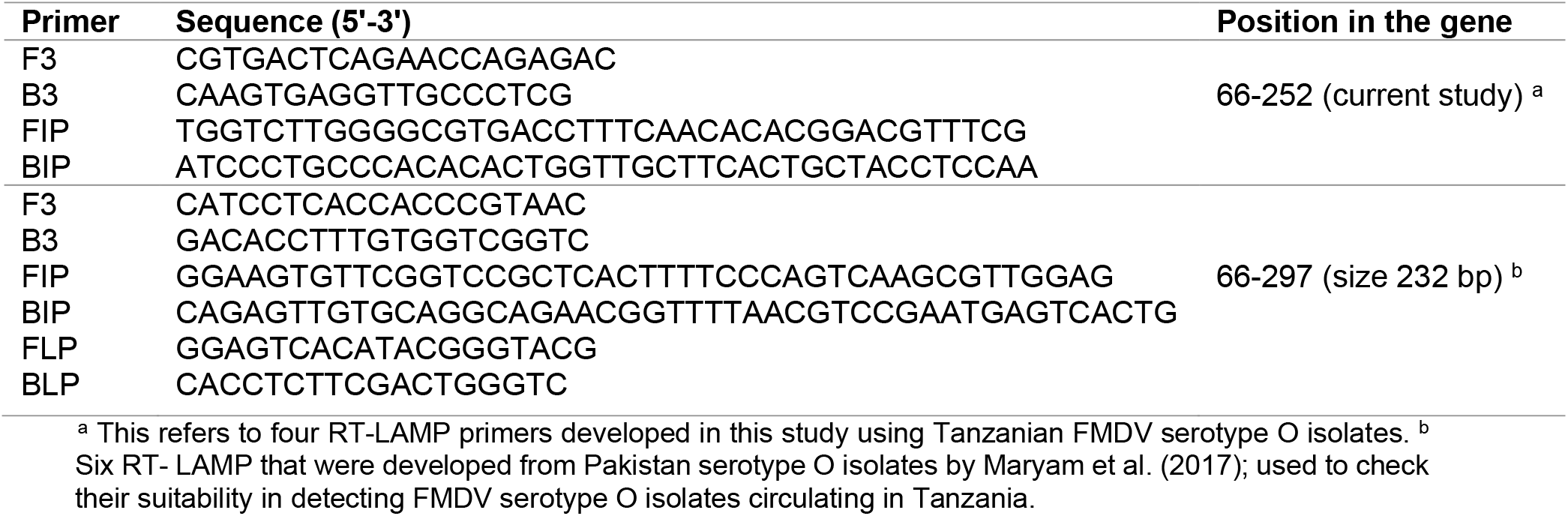
Details of RT-LAMP primers used for rapid serotyping of FMDV serotype O

**Table 3.**
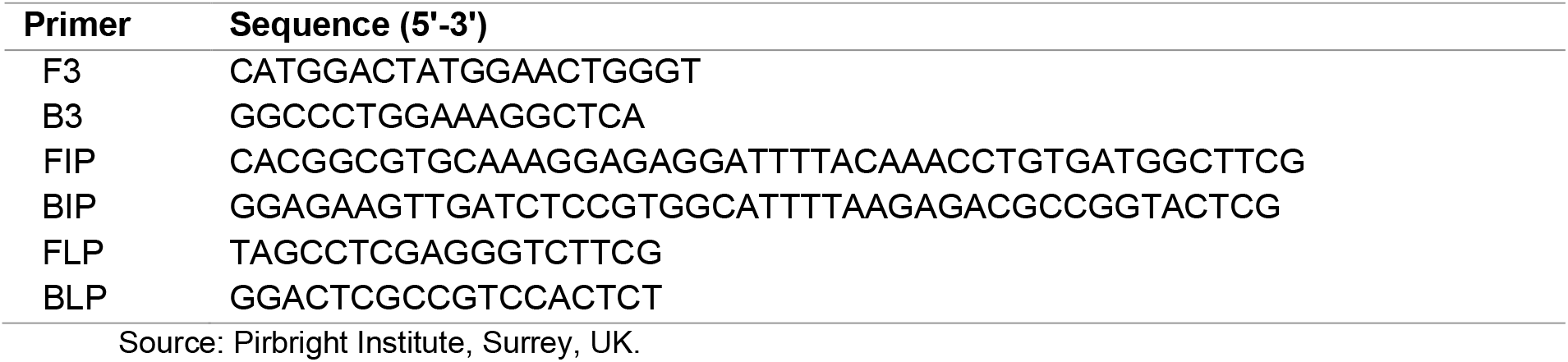
Details of RT-LAMP assay primer sets used for amplification of 3D polymerase gene of FMDV

**Table 4.** RT-LAMP assay results for detection and typing of serotype O FMDV

| Sample ID | Location (Tanzania) | Year | RT-LAMP (PAN) |  | RT-LAMP Serotyping |  | Results |
| --- | --- | --- | --- | --- | --- | --- | --- |
|  |  |  | T<sub>P</sub><sup>a</sup> (min:sec) | T<sub>A</sub><sup>b</sup> (°C) | T<sub>P</sub><sup>a</sup> (min:sec) | T<sub>A</sub><sup>b</sup> (°C) |  |
| 19 (PC)<sup>c</sup> | Fulwe, Morogoro | 2016 | 11.00 | 86.49 | 14.45 | 88.28 | + |
| 184 | Mtwara | 2017 | 17.00 | 87.39 | 15.30 | 87.78 | + |
| 254 | Kibaha, Rehema | 2017 | 10.15 | 86.99 | 13:45 | 87.89 | + |
| 257 | Kibaha, Rehema | 2017 | 16.00 | 86.19 | 15:00 | 87.88 | + |
| 259 | Kibaha, Rehema | 2017 | 16.00 | 86.92 | 15:45 | 87.88 | + |
| 262 | Kibaha, Rehema | 2017 | 10.30 | 87.41 | 10:45 | 88.28 | + |
| 265 | Kibaha, TC | 2017 | 13.45 | 86.79 | 18:30 | 88.68 | + |
| 268 | Kibaha, TC | 2017 | 10.15 | 87.38 | 16:30 | 88.08 | + |
| 272 | Kibaha, TC | 2017 | 13.00 | 86.10 | 25:45 | 87.02 | + |
| 307 | Kasulo, Ngara | 2017 | 13.00 | 86.79 | 16.00 | 87.39 | + |
| 365 | Tandahimba, Mtwara | 2017 | 11.15 | 87.11 | 22:30 | 88.11 | + |
| 369 | Tandahimba, Naputa | 2017 | 11.00 | 86.13 | 17.15 | 87.39 | + |
| 624 | Kinondoni | 2017 | 12.00 | 87.19 | 17:15 | 88.29 | + |
| 625 | Kinondoni | 2017 | 10.15 | 87.78 | 17:15 | 88.59 | + |

<sup>a</sup> Time to positivity (fluorescence emitted at one minute interval). <sup>b</sup> Annealing temperature. <sup>c</sup> Positive control

### 2.3. Design of Primers for RT-LAMP

Serotype O *VP1* sequences were used for designing RT-LAMP primers. Four primers, two outer (F3 and B3), and two inner (FIP and BIP), were designed using Primer Explorer V5 software (http://primerexplorer.jp/lampv5e/index.html, Fujitsu System Solutions, Ltd., Japan). The RT-LAMP primer amplify six distinct amplicons from the target sequence. The details of RT-LAMP primers are provided in Table 2.

### 2.4. RNA Extraction

Total RNA was extracted from the epithelial suspensions using RNeasy® Mini-kit (Qiagen, Hildan, Germany) as instructed by the manufacturer. RNA was also extracted from positive control (FMDV serotype O sample 19, previously tested and confirmed by RT-PCR and Sanger sequencing) and negative control (A non-template; nuclease free water). Viral RNA was extracted one from (Used in detection limit evaluation) using viral RNeasy® Mini-kit (Qiagen, Hildan, Germany). Extracted RNA was stored at −70°C until further use.

### 2.5. Amplification of 3D Polymerase Gene of Serotype O FMDV

To determine the presence of viral RNA, real time RT-LAMP (rRT-LAMP) assay was conducted using the primers listed in Table 3 following recommended incubation time of 30 min at 65°C on Genie® II (OptiGene Ltd, Horsham, UK). The reaction was terminated by incubating at 98°C for 1 min, followed by cooling to 80°C at a ramp rate of 0.05°C per second. Final reaction volume was 25ul constituting of; isothermal master mix (15μl), 5μl primer mixture [(F3/B3) 5 pmol: (FIP/BIP) 20 pmol: (FLP/BLP) 10 pmol] and 5μl extracted RNA template. Samples were defined as positive if both amplification and annealing of the LAMP product occurred. Amplified RT-LAMP products were visualized on a GelRed stained 1.5% agarose gel using E-Box imaging system (Vilber Lourmat).

### 2.6. RT-LAMP Assay Optimization

The RT-LAMP reaction conditions were optimized by performing test amplifications at varying reaction times (30, 45 and 60 min), temperatures (63°C, 64°C and 65°C), addition of avian myeloblastosis virus (AMV) reverse transcriptase enzyme and Pakistan loop primers (Maryam et al., 2017).The tests were repeated until optimum conditions (temperature and time) were secured. RT-LAMP assay was performed in the presence and absence of AMV reverse transcriptase enzyme. The reaction was terminated by incubation at 98°C for one min, followed by cooling to 80°C ramping at 0.05°C. The reaction mixture components were; isothermal master mix (15μl), 5μl primer mixture [(F3/B3) 5 pmol: (FIP/BIP) 20 pmol: (FLP/BLP) 10 pmol], AMV reverse transcriptase and 5μl extracted RNA template.

### 2.7. RT-LAMP Assay for Serotype O FMDV

The rRT-LAMP assay for serotype O FMDV was performed using the optimized conditions on Genie® II (OptiGene Ltd, Horsham, UK). Final reaction volume was 25μl constituting of; isothermal master mix (15μl), 5μl primer mixture [(F3/B3) 5 pmol: (FIP/BIP) 20 pmol: (FLP/BLP) 10 pmol], AMV reverse transcriptase enzyme and 5μl of extracted RNA template from 13 samples of serotype (table 4). The assay was performed at 65°C for 45 min, terminated by incubating at 98°C for 1 min, followed by cooling to 80°C at a ramp rate of 0.05°C per second on Genie ® II (OptiGene Ltd. Horsham, UK). Time to positivity (fluorescence emitted at one minute interval) and annealing temperature calculations were automated using Genie® Explorer v2.0.7.11 software (OptiGene Ltd.). Samples were considered positive if both amplification and annealing of the LAMP product occurred. The same protocol was used to amplify the *VP1* gene of serotype O FMDV using Pakistan RT-LAMP primers (table 2). Amplified RT-LAMP products were visualized on a GelRed stained 1.5% agarose gel using E-Box imaging system (Vilber Lourmat).

### 2.8. Specificity and Sensitivity of RT-LAMP

To assess the specificity of the FMDV serotype O RT-LAMP assay, cross-reactivity studies was conducted with RNA extracts from three FMDV Isolates of serotypes; A & SAT1 (Tanzania), SAT3 from Zambia and Namibia (Table 1). The cross-reactivity studies were performed using six sets of primers for serotypes; A, SAT 1 and SAT 3. Serotype O (sample ID: 19) was used as a positive control. Sensitivity of the RT-LAMP assay (i.e., detection limit) was assessed using ten-fold serial dilutions (10, 10^-2^, 10^-3^, 10^-4^, 10^-5^, 10^-6^, and 10^-7^) using one viral RNA template (sample ID: 19). The diluted viral RNA concentration was quantified using Epoch microplate spectrophotometer (Biotek, USA). The diluted viral RNA was used in RT-LAMP assay.

## 3. Results

### 3.1. RT-LAMP Amplification of 3D Polymerase Gene of FMDV (PAN)

The RT-LAMP assay detected viral RNA from 13 FMDV positive samples. The assay amplifies the 3D polymerase gene of serotype O at 65°C for 30 min, with time to positivity (T_P_) values and annealing temperature (T_A_) ranging between 10-17 min and 86.1-87.8°C respectively (Table 4).

Agarose gel electrophoresis of the RT-LAMP products (post-amplification analysis), results showed ladder-like bands (Figure 2)

**Figure 2.**
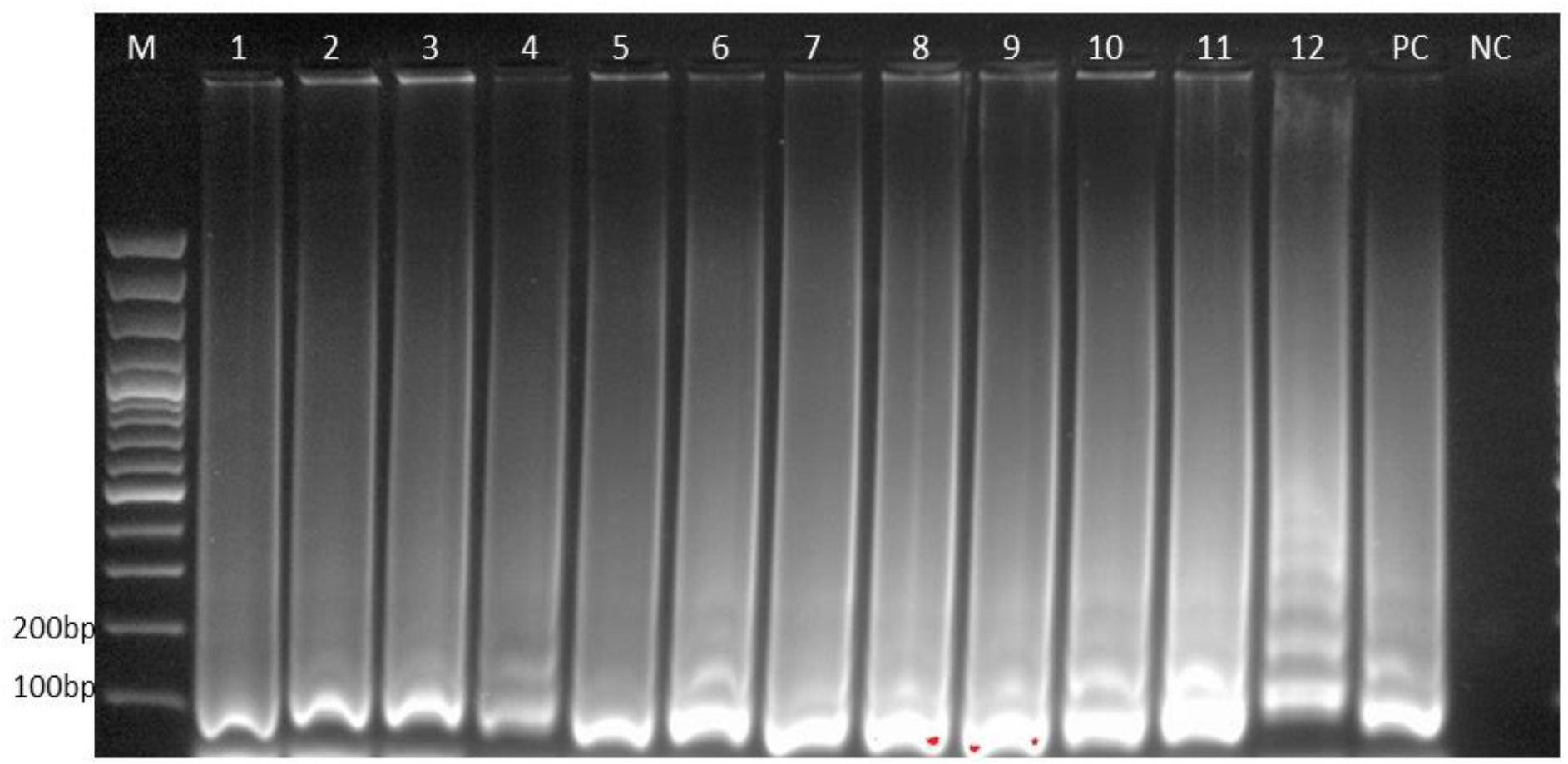
Agarose gel electrophoresis of FMD viral detection RT-LAMP assay products on a 1.5% agarose gel. Legend: lane M, 100 bp DNA ladder (New England BioLabs); lane 1-12, positive RT-LAMP reactions of test samples (table 4) as visualized by GelRed dye. Gel results show ladder-like bands, which confirms amplification of 3D polymerase gene, hence presence of FMD viral RNA; lane PC, positive control; lane NC, negative control (No template)

### 3.2. RT-LAMP Assay Optimization

The RT-LAMP assay successfully amplified the *VP1* gene of serotype O FMDV at 65°C for 45 minutes in presence of AMV reverse transcriptase enzyme and loop primers. The optimized RT-LAMP assay could reproduce the same results when repeated under the same conditions.

### 3.3. RT-LAMP Assay for FMDV Serotype O

The results of real time RT-LAMP assay for serotype O were assessed by time to positivity (T_P_) and annealing temperature (T_A_). Samples that showed both T_P_ and T_A_ were considered positive. RT-LAMP assay successfully amplified *VP1* gene of serotype O FMDV between 13 mins to 26 mins, and recorded T_A_ values between 70.0-89.0°C (Table 4). The amplification plot and annealing derivative plot (Figure 3) of the RT-LAMP reaction is shown.

**Figure 3.**
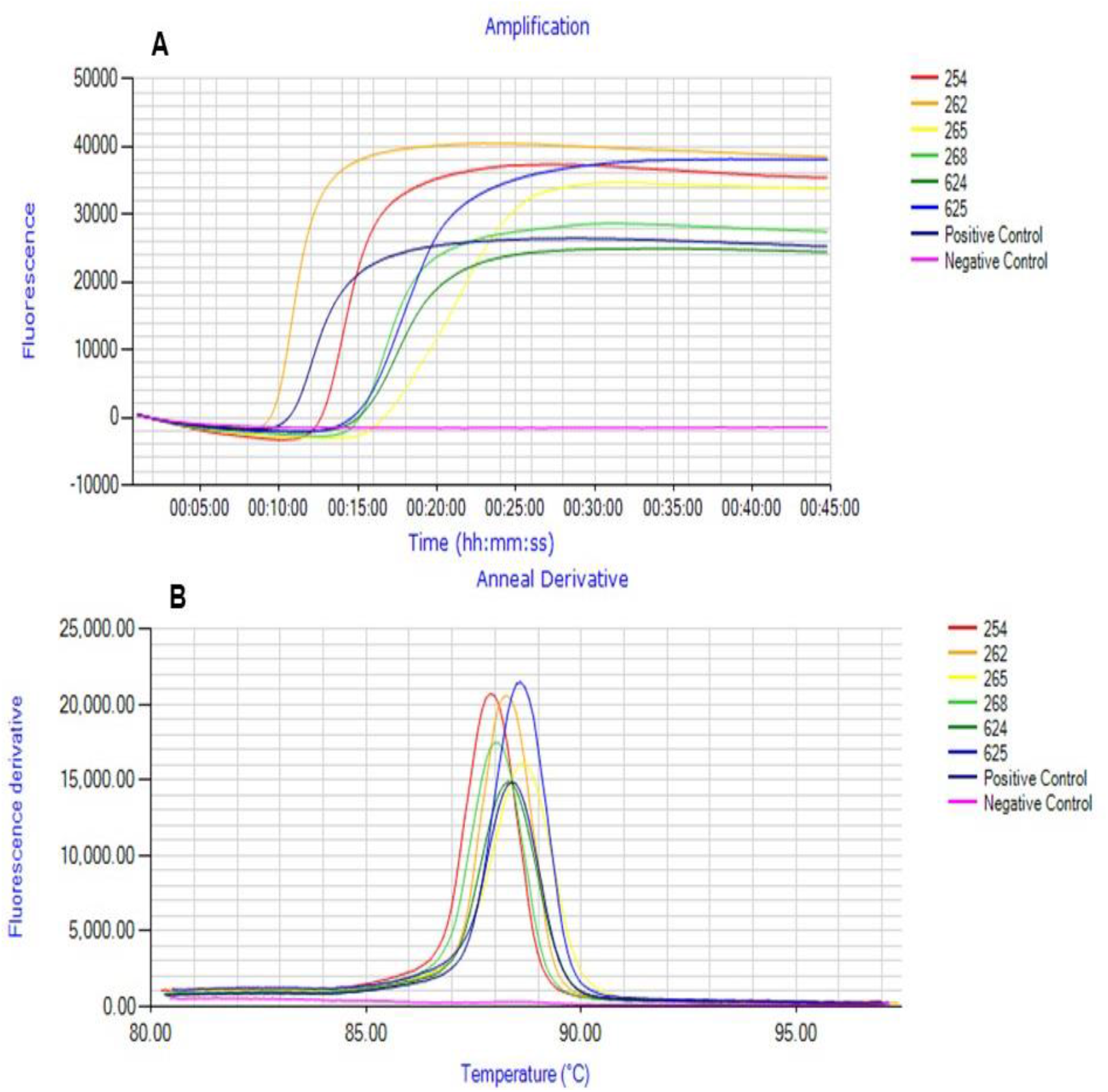
**(A)** Amplification plot and **(B)** annealing derivatives plot for FMDV serotype O obtained during RT-LAMP Assay. Legend: 254, 262, 265, 268, 624, and 625 are serotype O test samples used in the assay reaction. Other sample as indicated in (table 4) are not captured in these results (One run facilitates only 8 samples, inclusive of the controls). The generation of the amplification curves and the anneal derivatives indicates detection of *VP1* gene of serotype O FMDV by RT-LAMP, hence positive result.

This observation was consistent with agarose gel results where ladder-like bands were observed, indicating positive amplification of *VP1* gene (Figure 4). Amplification of *VP1* gene of serotype O FMDV isolates using Pakistan LAMP primers did not occur. However, when Pakistan loop primers developed by Maryam *et al*., (2017) were included in the assay, they amplified Tanzanian FMDV serotype O isolates as seen by a reduction in time to positivity.

**Figure 4.**
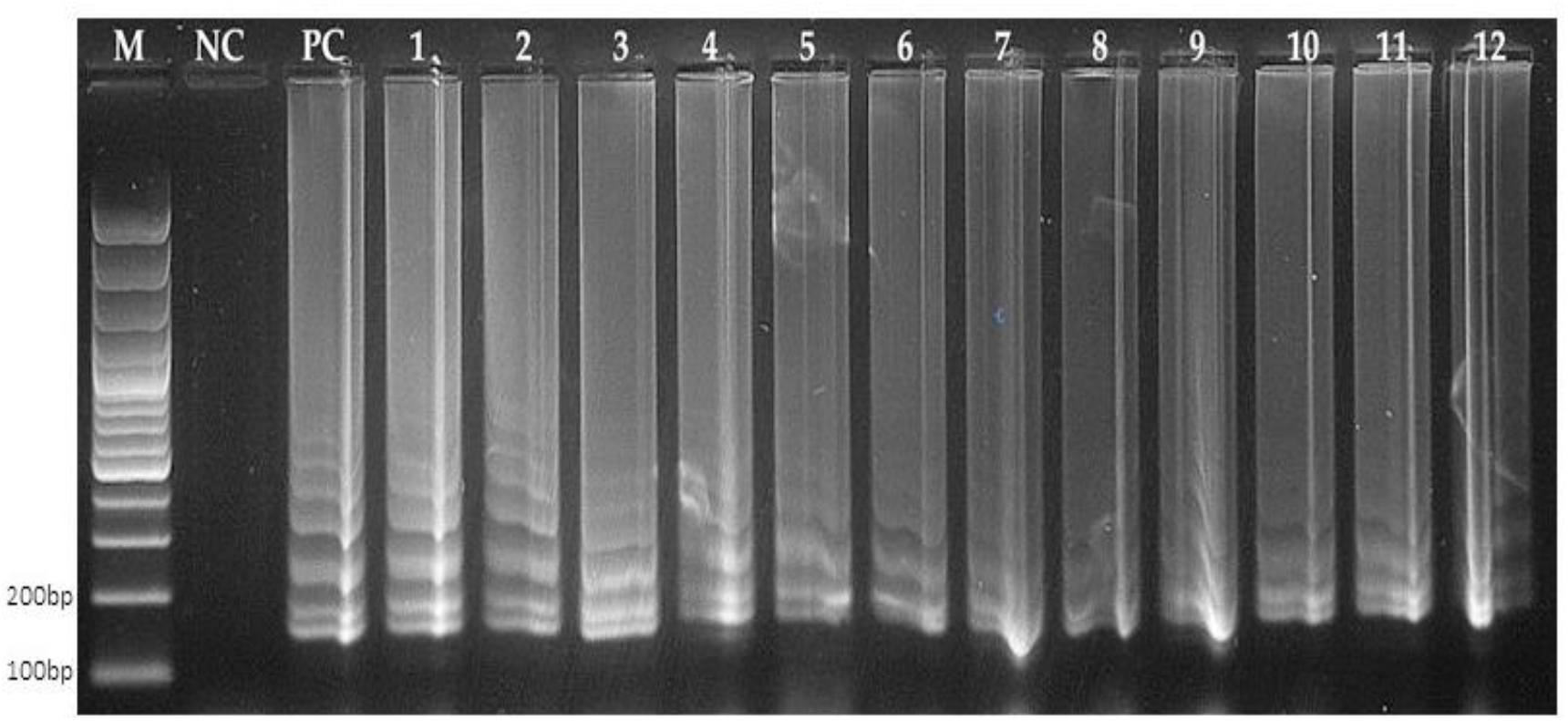
Agarose gel electrophoresis of serotype O FMDV RT-LAMP assay products on a 1.5% agarose gel. Legend: lane M, 100 bp DNA ladder (New England BioLabs); lane NC, negative control, no template; lane PC, positive control; lane 1-12, positive RT-LAMP reactions of test samples (table 4) as visualized by GelRed dye. Gel results show ladder-like bands, which confirms amplification of the serotype O *VP1* gene.

### 3.4. Specificity and Sensitivity of RT-LAMP Assay

The RT-LAMP primers for serotype O FMDV did not show any cross-reactivity with serotypes A, SAT 1 and SAT 3 (Figure 5). The assay was found to be specific for serotype O viruses.

**Figure 5.**
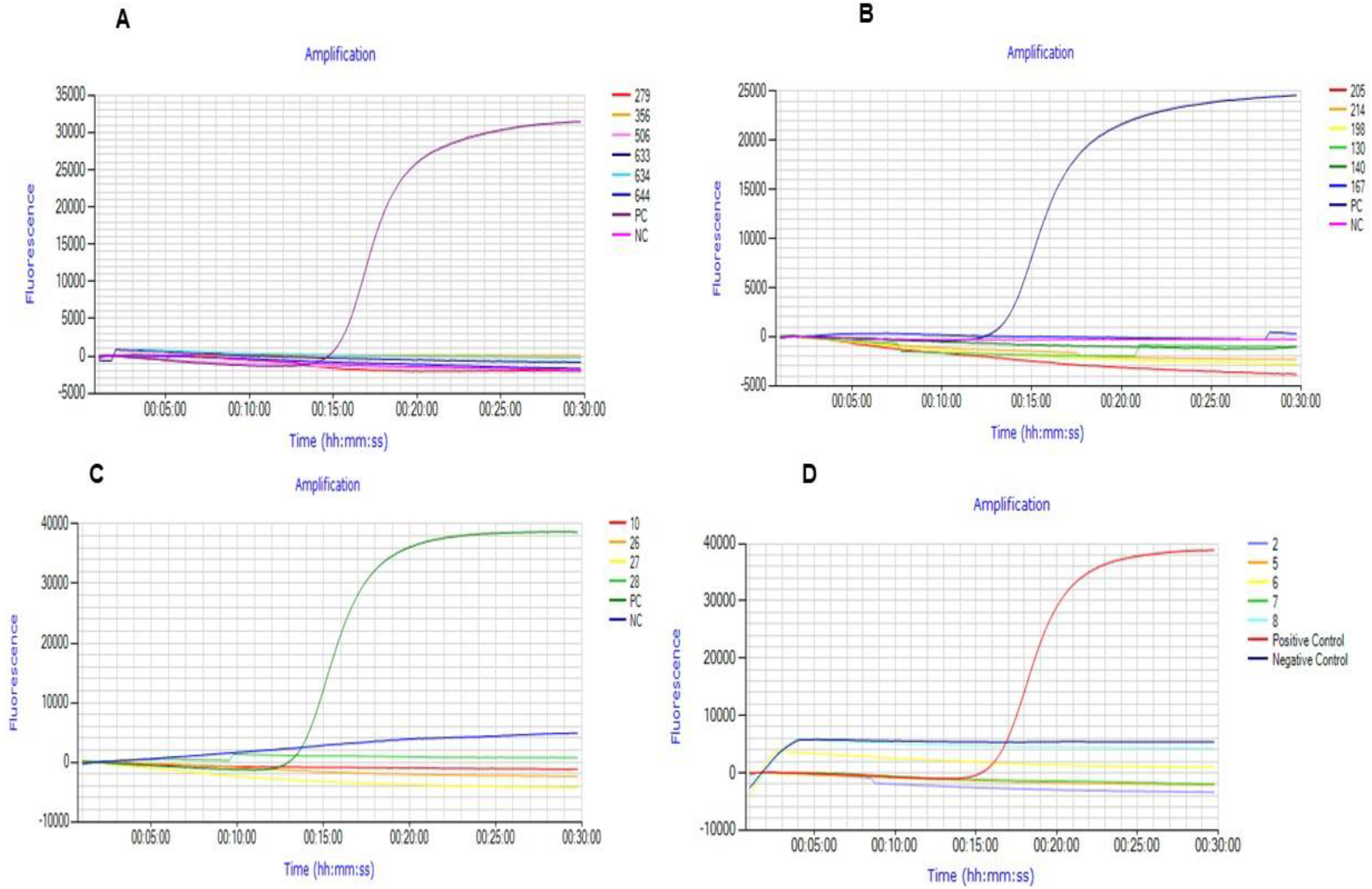
Amplification plots of RT-LAMP reaction of assessing specificity. Only the positive control (PC) FMDV serotype O, produced an amplification curve as shown in A (Pink), B (Purple), C (Green) and D (Red). **(A), (B), (C) and (D)** shows the amplification plots generated when the assay was tested against FMDV serotype; A, SAT 1, SAT 3 (Zambia) and SAT 3 (Namibia) respectively, which indicated no reaction occurred; Negative control (NC), No template control.

The detection limit of RT-LAMP assay was about 3.78 × 10^-2^ ng/μl at 10^-2^ dilution as shown in the amplification plot and gel results (Figure 6).

**Figure 6.**
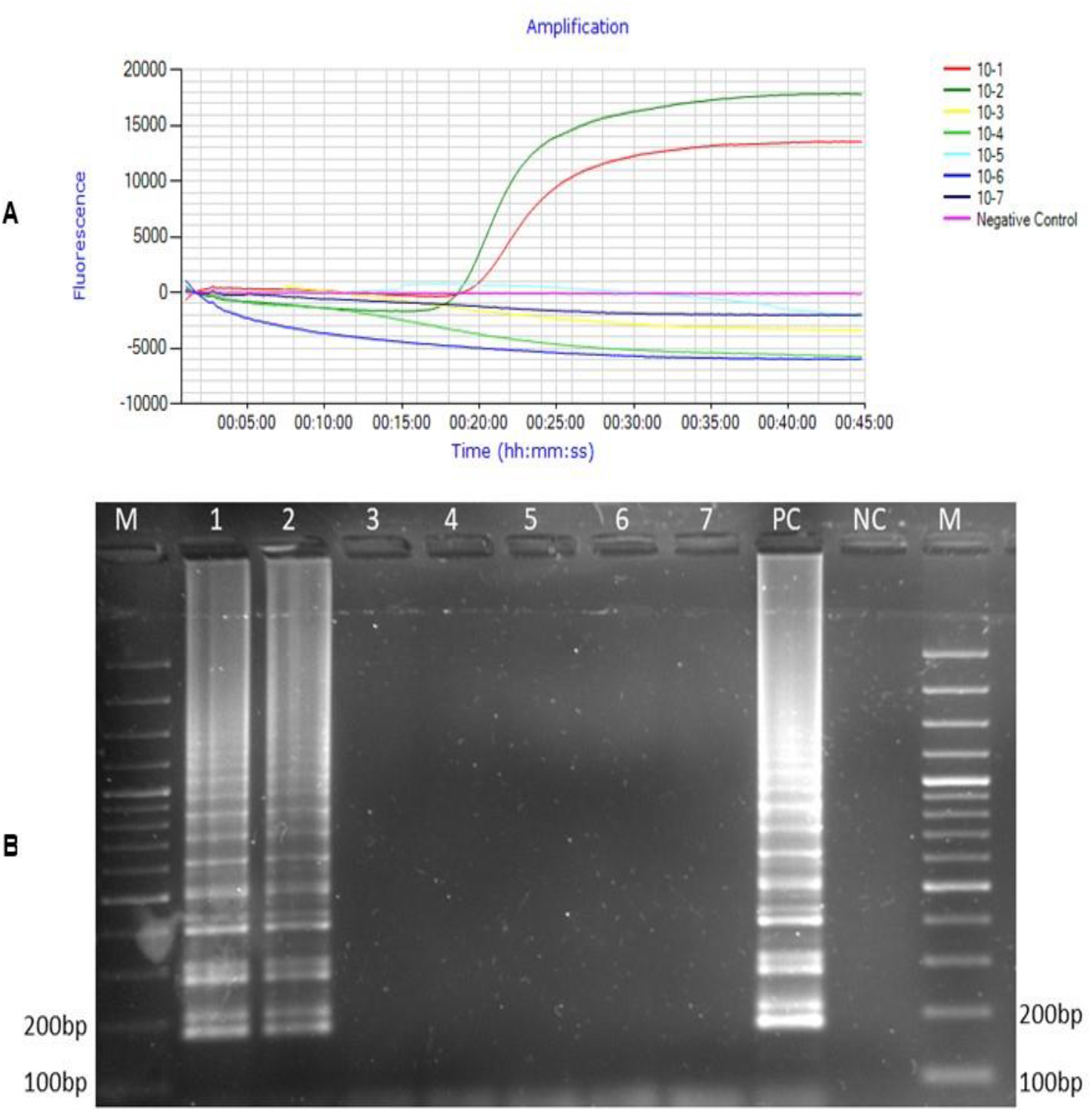
Assessing the sensitivity of the RT-LAMP assay. **(A)** Amplification plot for evaluation of detection limit of RT-LAMP assay for serotype O FMDV. As indicated by decreasing concentrations of viral RNA (10-1 to 10-7 dilutions per test tube), the assay amplified serotype O up to 10-2 dilution with viral concentration of 3.78 ×10-2 ng/μl. **(B)** Agarose gel electrophoresis of detection limit RT-LAMP assay products on a 1.5% agarose gel; Legend: lane M, 100 bp DNA ladder (New England BioLabs); lane 1-7 (Viral RNA decreasing concentrations of 10-1 to 10-7); lane PC, positive control; lane NC, negative control. As shown in the figure, the RT-LAMP assay detected the VP1 gene of serotype O up to 10-2 dilution having a concentration of 3.78 ×10-2 ng/μl viral RNA.

## 4. Discussion

Rapid detection of FMDV serotype O is vital in the surveillance, management, and control of FMD. The aim of this study was to develop RT-LAMP primers suitable for detecting *VP1* gene of serotype O FMDV strains circulating in Tanzania as well as to check the suitability of Pakistan RT-LAMP primers on the same.

The findings of this study indicated that all samples (n=13) tested positive for FMD viral RNA as indicated by both the presence of time to positivity and annealing temperature. This observation was consistent with the results of agarose gel electrophoresis, where ladder-like bands were observed, indicating amplified positive product. RT-LAMP assay was able to detect FMD viral RNA between 10-17 mins, results which agree with previous findings (Dukes et al., 2006; Kasanga et al., 2014) which demonstrated the ability of RT-LAMP assay to detect FMDV within 60 mins.

The RT-LAMP primers developed in this study successfully amplified the *VP1* gene of serotype O FMDV contrary to those developed by (Maryam et al., 2017), which failed to amplify serotype O virus isolates recovered from cattle in Tanzania. The failure of these primers to amplify the *VP1* polyprotein gene from the Tanzania serotype O viruses could be ascribed to genetic diversity which classify the Pakistan serotype O viruses under different pool and genotype (Cathay and Middle East-South Asia - ME-SA) as compared to those from Tanzania (EA-2) (Knowles & Samuel, 2003).The assay was able to detect the *VP1* gene at 13-26 mins in the presence of AMV reverse transcriptase and loop primers but for optimum reaction, assay reaction time was set for 45 mins. These results agree with previous RT-LAMP assay findings by (Farooq et al., 2015; Lim et al., 2018; Madhanmohan et al., 2013; Maryam et al., 2017) that demonstrated RT-LAMP to detect *VP1* gene of serotype O viruses within 60 mins. The RT-LAMP annealing temperature (Derivative) ranged between 87.0°C-89.0°C with a min of 87.02 ± 0.3 °C.

The inclusion of loop primers in the assay had significant impact on the amplification as seen in a reduction of time to positivity compared to a reaction without loop primers. These results agree with previous studies conducted by (Madhanmohan et al., 2013; Maryam et al., 2017; Mori & Notomi, 2009). It was observed that loop primers developed by (Maryam et al., 2017) amplified the *VP1* gene of serotype O virus isolates circulating in Tanzania. This finding deduced that, these loop primers targeted a conserved region within the *VP1* gene as they were able to successfully amplify both serotype O virus isolates from Pakistan and Tanzania. It was also noted that addition of AMV reverse transcriptase had significant impact on amplification rate as seen by the reduction of time to positivity (TP).

The efficiency of RT-LAMP assay relies heavily on the specificity of the primer sets designed (Madhanmohan et al., 2013). RT-LAMP primers were specific to serotype O viruses as no cross-reactivity with other serotypes was evidenced. These results agree with previous studies conducted by (Madhanmohan et al., 2013; Maryam et al., 2017). The sensitivity of RT-LAMP assay was inferred from the detection limit findings, which was about 3.78 ×10-2 ng/μl (3.78 ×10-3 μg/μl) and consistent gel results. This assay is able to detect low copies of viral RNA in FMD suspected samples, findings similar to that made by (Madhanmohan et al., 2013).

## 5. Conclusions

This molecular diagnostic approach has potential future application in improving FMDV surveillance as it provides baseline information for controlling foot-and-mouth disease (FMD) outbreaks in Tanzania. The study recommends further evaluation of the assay to determine whether it is feasible in detecting serotype O viruses circulating in other regions in Tanzania. It also recommends development of RT-LAMP assay for detecting other circulating serotypes A, SAT1 and SAT 2.

## Acknowledgments

The authors are thankful to Professor Gerald Misinzo, Dr. Henry Muriuki, Dr. Jean Hakizimana and Ms. Juliana for guidance and technical support during the study. The actors acknowledge the Intermediate Fellowship in Public Health and Tropical Medicine (IFPHTM) project for FMDV surveillance, in the College of Veterinary Medicine and Biomedical Sciences, Sokoine University of Agriculture, Tanzania for providing the samples used this study.

